# Stranger Swings: Temperature-Dependent Upsides and Downsides of a Densovirus in *Aedes albopictus*

**DOI:** 10.64898/2026.03.09.710552

**Authors:** C Boëte, M Perriat-Sanguinet, A-S Gosselin-Grenet, P Makoundou, M Ogliastro, M Sicard, S Unal, M Weill, C Atyame

**Affiliations:** ISEM, Univ Montpellier, CNRS, IRD, EPHE, Montpellier, France; DGIMI, Univ Montpellier, INRAE, Montpellier, France; Université de La Réunion, UMR PIMIT (Processus Infectieux en Milieu Insulaire Tropical) CNRS 9192, INSERM 1187, IRD 249, Saint-Denis, île de La Réunion, France

## Abstract

Vector-borne diseases remain a significant global health concern, with the invasive mosquito *Aedes albopictus* playing a key role in the transmission of arboviruses including dengue, chikungunya, and Zika viruses. As this species expands in novel territories, effective vector control strategies are increasingly critical. Densoviruses (DVs) have emerged as potential biological control agents, either through direct pathogenic effects on mosquito populations or via paratransgenesis. However, the influence of combined environmental factors, such as temperature and densovirus infection on mosquito life-history traits remains largely unexplored.

In this study, we investigated the effects of different temperatures (28°C, 31°C, and 34°C) and exposure to the densovirus AalDV2 on the survival of *Ae. albopictus* and on several mosquito life-history traits including its development time, size and symmetry at adult stage. Larvae were individually reared under controlled conditions and exposed to AalDV2 or a control treatment.

Temperature had a strong nonlinear effect on survival, with dramatic mortality increases at 34°C. Unexpectedly, AalDV2-infected larvae showed significantly higher survival than controls at this extreme temperature, suggesting a protective effect under thermal stress.

Across all temperatures, viral infection delayed pupation in a sex-dependent manner, with females experiencing greater delays and reduced adult wing size. Quantitative PCR revealed high infection rates (>96%) across all conditions with a viral load increasing with higher temperature in a sex-dependent manner.

Our findings reveal a paradoxical outcome: while AalDV2 imposes fitness costs through delayed development and reduced body size, it confers an unexpected survival advantage at the extreme temperature. This represents the first documentation of temperature-dependent protective effects by an entomopathogenic virus in mosquitoes, challenging conventional assumptions about pathogen impacts. These results have critical implications for vector biocontrol strategies in a warming climate, as densovirus deployment could inadvertently enhance mosquito resilience in heat-stressed regions. Further research is needed to elucidate the underlying mechanisms and assess the impact on mosquito population dynamics.

## Introduction

Vector-borne diseases continue to pose a serious threat worldwide and the last decades have seen the emergence of Chikungunya and Zika viruses as well as multiple large-scale dengue outbreaks. These *Aedes*-transmitted neglected tropical diseases are highly sensitive to environmental factors including temperature (Lambrechts et al. 2011, Mordecai et al. 2017, Delrieu et al. 2023) shaping mosquito survival and vectorial capacity. Consequently, environmental variability plays a critical role in determining the timing, intensity, and geographic distribution of disease transmission, highlighting the importance of climate-informed strategies not only for predicting and controlling outbreaks but also for optimizing vector control efforts. Compounding the challenge, one of the major vectors of arboviruses, *Aedes albopictus*, is an invasive species with great adaptability to a diversity of ecological conditions and it is expanding its geographic distribution (Ryan et al. 2019) not only in tropical but also in temperate regions (Farooq et al. 2025). Although vaccines exist for some of these viruses, their use is restricted to specific population groups, leaving critical gaps in coverage that prevent achieving herd immunity and effective population-level protection. This leads to vector control being the major way to fight against those diseases (Kraemer et al. 2019, de Souza and Weaver 2024, Abbasi 2025). When considering the arboviral outbreaks that occurred in the last decades, a number of responses have been brought ranging from vaccination against Yellow Fever in Angola in 2016 (Boëte 2016) to the removal of breeding site, the use of larvicide and the killing of adult via fumigation in various contexts (Dengue in La Réunion, 2018; Zika in Brazil, 2015) with inherent limitations due to the insecticide resistance in vectors. While some methods are well-known for their efficacy when properly deployed, the difficulties in the management of epidemics leads to the need to consider novel approaches to combat vector-borne diseases (Boëte 2021). Aside from the development of innovative approaches, lines of research on entomopathogenic densoviruses (DV), small DNA viruses belonging to the *Parvoviridae* family, have been showing some potential for biological mosquito control decades ago (Buchatskiĭ et *al.* 1987, Kuznetsova and Buchatskiĭ 1988) and have been revisited in the recent years (Johnson and Rasgon 2018; Perrin et *al.* 2020). Densoviruses are structurally simple, non-enveloped viruses with a single-stranded DNA genome, known for their diverse host range among arthropods, including mosquitoes, lepidopterans, crustaceans, and even echinoderms. Two approaches regarding their potential for vector control are nowadays considered: the use of their pathogenicity to reduce mosquito population or, on the contrary, their low impact on survival and fecundity that could make them interesting candidate for paratransgenesis aiming at modifying population of vectors. Despite several studies on the pathogenicity of DVs to different mosquito species (Ledermann et *al.* 2004, Carlson et *al.* 2006) or strains (Perrin et *al.* 2020), there are still a number of obstacles before any potential development and deployment of such a tool to be considered in integrated vector management programs (Berger et *al.* 2025). Among them the impact of temperature, known to be one of the major parameters in the interactions between mosquitoes and their infecting viruses (Murdock et *al.* 2012, Carlassara et *al.* 2024, Terradas et *al.* 2024), on both the outcome of the infection by DVs and its impact on the life-history traits of mosquitoes is still highly unexplored (Li et *al.* 2019). Our study aims then at addressing this point by measuring the life-history traits of *Ae. albopictus* infected by the densovirus AalDV2 under different temperatures ranging from 28°C to 34°C.

## Methods

### Virus production and quantification

The AalDV2 densovirus strain used in this study was isolated from a chronically infected cell line of the C6/36 clone (Jousset et *al.* 1993), which originated from homogenized *A. albopictus* larvae. It was formerly known as AaPV for Aedes albopictus parvovirus and was found to be pathogenic for *Ae. aegypti* and *Ae. albopictus*. This cell line was obtained from the Institut Pasteur in Paris. C6/36 cells were cultured at 28°C in RPMI cell culture medium (Gibco), supplemented with 10% heat-inactivated fetal bovine serum (FBS; Gibco), 1% antibiotic-antifungal cocktail (ATB, Gibco), and 1% non-essential amino acids (NEAA; Gibco). The purification of AalDV2 densovirus from C6/36 cells involves an initial step of cell lysis through successive freeze-thaw cycles. The cells and their supernatants are pooled and centrifuged at 4°C for 15 minutes at 3000 g to separate large cellular debris from the virus. The supernatant is then layered onto a 15% sucrose cushion and subjected to ultracentrifugation at 38,000 revolutions per minute (SW41 rotor, Optima™-90K Beckman Ultracentrifuge) for 3 hours at 8°C. The resulting pellet is resuspended in a 1 mM Tris and 0.1 mM EDTA buffer (TE 0.1X). The final viral suspension was analyzed qualitatively by transmission electron microscopy and quantified by quantitative PCR (qPCR) in genome equivalent virus (gev) per µl as described in Perrin et *al.* (2020). Control (mock) productions were prepared in the same conditions from uninfected C6/36 cells.

### Larval and adult maintenance, infection protocol and temperature regimes

*Aedes albopictus* mosquitoes are originating from a colony established on Réunion Island and are maintained at hundreds of individuals per generation at ISEM, Montpellier, France. One day after hatching, mosquito larvae were individually placed in 96 well-plates with 180 µL of deionized water at one of the following conditions 28±1°C, 31±1°C and 34±1°C. For mosquitoes exposed to the AaDV2, 10^10^ virus equivalent genome (veg) of purified virus (a dose similar to the one used in previous experiments (Perrin et *al.* 2020)) were added to in the water of each larva while for the control one, the same volume (30 µL) of an extract of an uninfected C6/36 culture was added and the final volume for each larva was 210 µL. All mosquitoes had the same food regime of Tetramin® baby fish food until pupation. On day 1 after hatching, larvae were given 0.04 µg, 0.06 on day 2, 0.12 on day 3, 0.24 on day 4 and 0.48 of all subsequent even days. On day 3, to ensure sufficient space for the larval development, all surviving larvae were transferred into a 12 well-plate with 3 mL of deionized water per well. Mosquitoes were reared in a 12:12 light/dark cycle, at 75 ±5% relative humidity. Larvae were checked every day to count the number of dead and live ones and to record age at pupation. Once individuals pupated, they were transferred into hemolysis tube with 1 mL of deionized water and, 24h after emergence, adults were sexed and frozen and their wing length, a proxy of their size and fecundity, was measured. Fluctuation asymmetry, representing the deviation from the perfect bilateral symmetry of wings has also been measured in order to evaluate the impact of the treatments (temperature and AalDV2 virus exposure) on the development stability. As each larva was reared individually, each mosquito is considered an individual point. The experiment was divided in 3 blocs with 2 different temperature conditions tested in each one leading to 2 independent replicates for each combination of temperature and infection treatment. Because mortality rate was expected to vary across subgroups, sample size was adjusted to ensure sufficient statistical power and representation in each group. Thus, in Bloc 1, the temperatures used were 28 ± 1°C and 31 ± 1°C; in Bloc 2, 28 ± 1°C and 34 ± 1°C were applied; while in Bloc 3, the temperatures were 31 ± 1°C and 34 ± 1°C.

### Virus quantification in mosquitoes: Measure of the infection prevalence and viral load

Real-time quantitative PCR was run using the LightCycler 480 system (Roche) and the SensiFAST SYBR No-ROX Kit (Meridian Biosciences). All DNA samples were obtained via phenol chloroform extraction and they were analyzed in triplicates for each quantification. The viral load (*i.e.*, the number of copies of AalDV2 per mosquito) was assessed with qPCR using a specific AalDV2 PCR normalized on the host *Ae. albopictus* specific single-copy actin locus. For AalDV2 and the actin the following protocol was used: 5 min at 95°C, 30 cycles of 94°C for 30 s, 52°C for 30 s, and 72°C for 60 s, then 72°C for 5 min. The primers for the amplification of the *Ae. albopictus’* actine primers were actAlb-dir: 5’-GCAAACGTGGTATCCTGAC-3’and actAlb-rev: 5’-GTCAGGAGAACTGGGTGCT-3’ (Tortosa et *al.* 2008) and for AalDV2 they were qAalDV2_AP_2_F: 5’-CTCTGGAGCCGCTGTGTAAT-3’ and qAalDV2_AP_2_R:5’ TGGCCAACAATTACGAACAA-3’ (Perrin et *al.* 2020). The ratio between the 2 signals was used to estimate the relative number of AalDV2 genomes per mosquito genome. When the level of amplification of actine, *i.e.* the cycle threshold, was below 25, samples were discarded.

### Statistical methods

All statistical analyses were conducted with RStudio version 2024.04.2+764. Data manipulation and preparation were performed using the *dplyr* package. For data visualization, the *ggplot2* package was employed. To ensure normality of residuals, data transformations were applied using the *bestNormalize* package, with the *orderNorm* transformation selected for variables such as age at pupation, wing size, and fluctuating asymmetry. For modeling, the lme4 package was used to fit linear mixed-effects models, while generalized linear mixed models were fitted using the *glmmTMB* package. The *lmerTest* package was used for the calculation of p-values for mixed models. To test the significance of the fixed effects and their interactions, a Type II or a Type III Wald chi-square test was conducted using the Anova function from the *car* package. For model selection and averaging, the *AICcmodavg* package was used. Post-hoc comparisons were conducted with *emmeans*, with p-value adjustments via *multcomp*. The verifications of the model assumptions were conducted with the *DHARMa* package, this includes simulations of residuals to test for uniformity (Kolmogorov-Smirnov) and dispersion issues.

Mortality at the aquatic stages (larvae and pupae) was analysed using generalized linear mixed models (GLMMs) with a binomial error distribution and the explanatory variables were temperature and exposure to AalDV2. Model robustness was assessed by comparing alternative model formulations, including heteroscedastic binomial models and a biologically motivated thermal threshold parameterization. The sex-ratio was analysed with a generalized linear model with a binomial distribution of errors where and the explanatory variables were the age at pupation, temperature and exposure to AalDV2.

After transformation the age at pupation, the wing length and their asymmetry were analyzed with a linear model with a normal distribution of errors and the explanatory variables were temperature, infection status and sex. The level of infection for the exposed mosquitoes was analyzed with a linear model with robust standard errors. The analysis was conducted using the *lm* function, followed by robust inference with the *coeftest* function from the *lmtest* package and vcovHC from the *sandwich* package to account for potential heteroscedasticity.

## Results

### Effect of temperature and exposure to AalDV2 on the survival of the aquatic stages

Across replicates, the survival of the aquatic stages remains above 88.4% at 28°C or 31°C (Table S1) regardless of exposure to AalDV2. In contrast it dramatically drops when the temperature reaches 34°C, revealing a significant negative impact of the temperature on the survival of the aquatic stages (χ² =280.74, df = 2, p < 2.2e-16) (Table 1; Fig. 1). This pattern indicates a nonlinear response of the mosquito to the temperature increase. This suggests a threshold-like effect, where mortality accelerates sharply beyond a critical temperature point. Consistent with this interpretation, our analysis confirmed a highly significant effect of temperature on survival, as well as a significant overall effect of AalDV2 exposure (χ² = 6.39, df = 1, p = 0.011). However, the interaction between temperature and viral exposure was not statistically significant (χ² = 3.51, df = 2, p = 0.17), indicating that treatment effects did not vary linearly across temperatures.

**Fig. 1:**
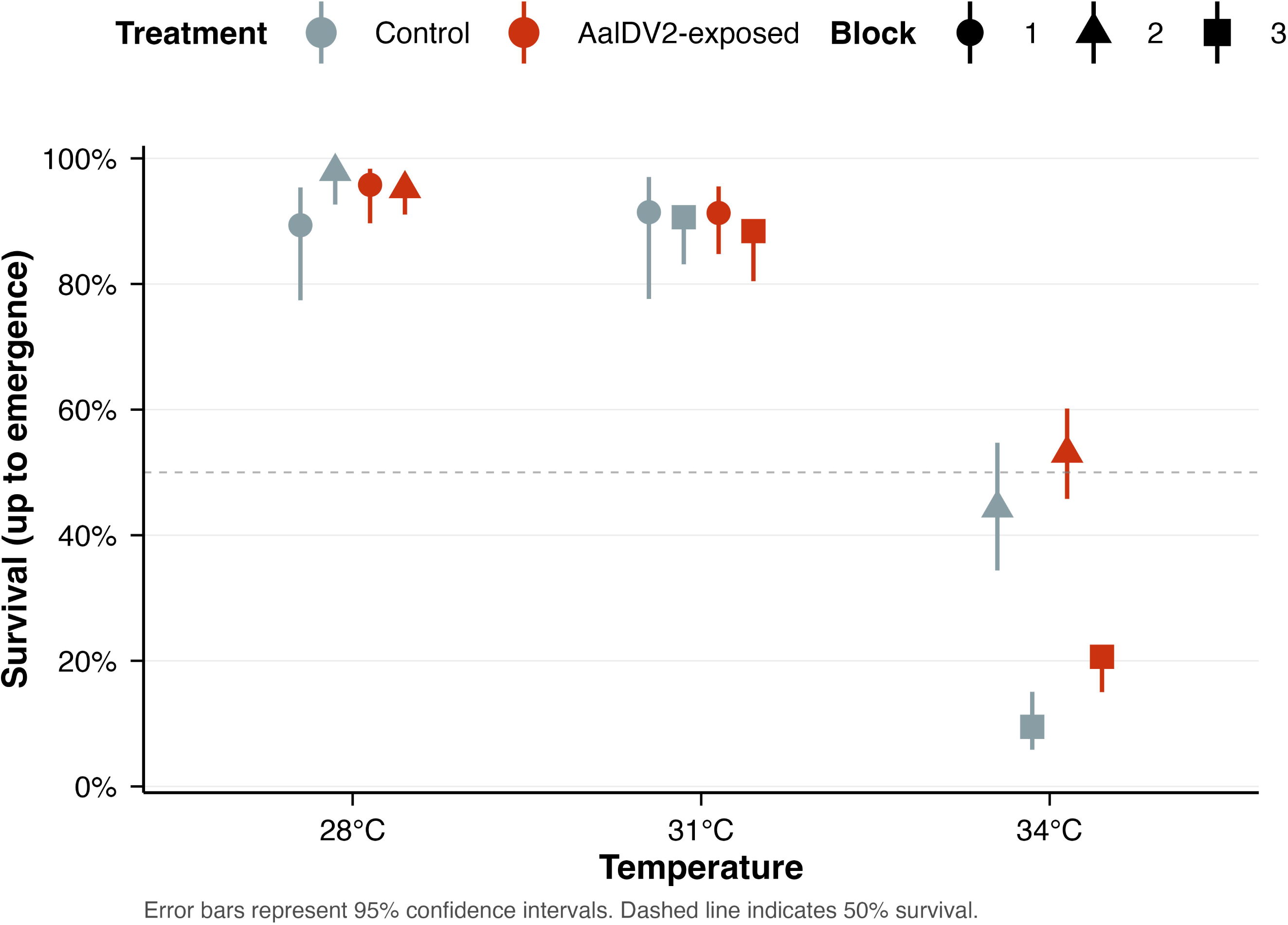
Effect of the exposure to the densovirus AalDV2 and of different temperatures (28°C, 31°C, and 34°C) on the survival of the aquatic stages (larvae and pupae) of the mosquito *Aedes albopictus*. Points represent the mean survival proportion, and error bars indicate 95% confidence intervals. Different shapes represent different blocks. The dashed line indicates a 50% survival threshold.

**Table 1:** Statistical analysis of the effect the temperature and the exposure to AalDV2 on the survival of *Aedes albopictus* larvae. The table presents chi-square (χ²) test results assessing the effect of AalDV2-exposure, temperature, and their interaction on larval survival. Degrees of freedom (df) and p-values (Pr(>χ²)) are provided for each factor. The number of asterisk*s* indicates the statistical significance levels as follows: *** (p < 0.001) and * (p < 0.05).

| Effect | $\chi^2$ | df | Pr(>$\chi^2$) |
| --- | --- | --- | --- |
| AalDV2- Exposure | 6.39 | 1 | 0.011 * |
| Temperature | 280.74 | 2 | < 2.2e-16 *** |
| Temperature : AalDV2-Exposure | 3.51 | 2 | 0.17 |

To assess whether the effect of AalDV2 exposure emerged specifically under thermal stress, we conducted post-hoc contrasts within each temperature level. No significant differences in survival were detected between AalDV2-exposed and unexposed larvae at 28°C or 31°C. In contrast, at 34°C, exposure to AalDV2 was associated with a significantly higher probability of survival (odds ratio = 1.87, p = 0.0018). At this temperature, survival to adulthood reached 27.69% and 53.04% in AalDV2-exposed larvae across the two replicates, compared with 9.49% and 44.32% in control larvae (Fig. 1).

Given the nonlinear temperature–mortality relationship, we further tested a thermal threshold model distinguishing normal (28–31°C) from high (34°C) thermal conditions. This model provided a fit comparable to the initial model while using fewer parameters. Under normal thermal conditions, AalDV2 exposure had no detectable effect on survival, whereas under high thermal stress, viral exposure significantly reduced mortality, confirming that the protective effect of AalDV2 emerges only once a critical thermal threshold is exceeded. A supplementary analysis stratifying aquatic mortality by developmental stage revealed distinct stage-specific patterns. At the larval stage, mortality was temperature-dependent but independent of AalDV2 exposure across all temperatures (28°C: odds ratio = 0.278, p = 0.234; 31°C: odds ratio = 0.679, p = 0.428; 34°C: odds ratio = 1.202, p = 0.299). In contrast, AalDV2 conferred significant protection specifically at the pupal stage, but only at the highest thermal stress level (34°C: odds ratio = 2.35, p = 0.0006), where pupal mortality in individuals unexposed to AalDV2 reached 56.1% compared with 33.8% in AalDV2-exposed individuals. At the lower temperatures, pupal mortality remained minimal (<5%) in both treatment groups with no significant differences (28°C: odds ratio = 2.01, p = 0.235; 31°C: odds ratio = 1.57, p = 0.510). These findings establish the pupal stage as the primary thermal sensitivity bottleneck, and reveal that AalDV2 provides condition-dependent protection exclusively during this critical developmental window under severe thermal stress.

Together, these results indicate that while AalDV2 exposure does not prevent temperature-induced mortality, it partially mitigates its impact under extreme thermal conditions, particularly at the pupal stage. This reveals a context-dependent protective effect of the virus at the upper limit of thermal tolerance of the mosquito aquatic stages.

### Impact on the sex-ratio

For all the different temperatures and whether mosquitoes were exposed to AalDV2 or not, the proportion of females increased with later pupation, a phenomenon usually observed in *Aedes* spp (Clements, 1992). Our study revealed no significant differences between infected and control groups (Table S2), although there was a trend towards a higher female-biased sex-ratio at 34°C in the infected groups (Fig. 2). While this was not statistically significant, it suggests that viral infection may slightly influence sex-ratio under stressful environmental conditions, as reported in previous studies (Perrin et *al.* 2020) while others have observed a much higher mortality rate in adult males infected at the larval stage with densoviruses (Buchatsky 1989).

**Fig. 2:**
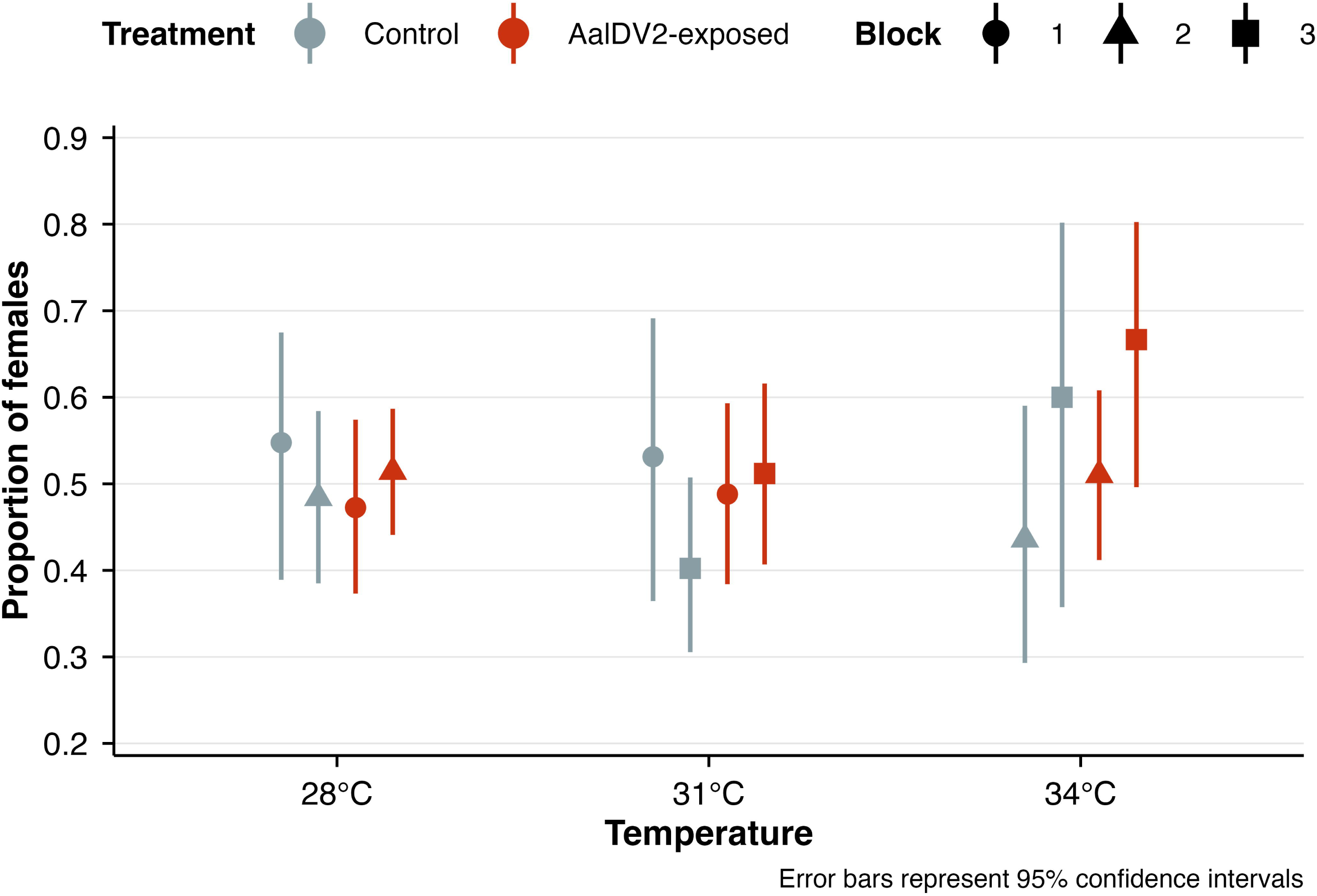
Impact of the exposure to the densovirus AalDV2 and of different temperatures (28°C, 31°C, and 34°C) on the sex-ratio of *Aedes albopictus* mosquitoes. Points represent the mean proportion of females, and error bars indicate 95% confidence intervals. Different shapes represent different blocks.

### Effect of temperature and exposure to AalDV2 on the life-history traits of mosquitoes

#### Impact on the age at pupation

The analysis of the age at pupation (Table 2) was conducted on 872 individuals that survived to adulthood. The model revealed significant effects of temperature (χ² = 739.66, df = 2, *p* < 2.2e-16), sex (χ² = 147.42, df = 1, *p* < 2.2e-16), and AalDV2- exposure (χ² = 4.37, df = 1, *p* = 0.0366). Additionally, there was a significant interaction between AalDV2-exposure and sex (χ² = 3.92, df = 1, *p* = 0.0477) (Fig. 3). It appears indeed that a higher temperature leads to a longer time to reach pupation and that, in any condition, males pupate earlier than females as previously described (Wormington and Juliano 2014, Teder et *al.* 2021). Regarding the exposure to AalDV2, it has a significant effect on the age at pupation, with a stronger delay observed in females compared to males, as evidenced by the significant interaction between sex and AalDV2-exposure (*p* = 0.0477).

**Fig. 3.**
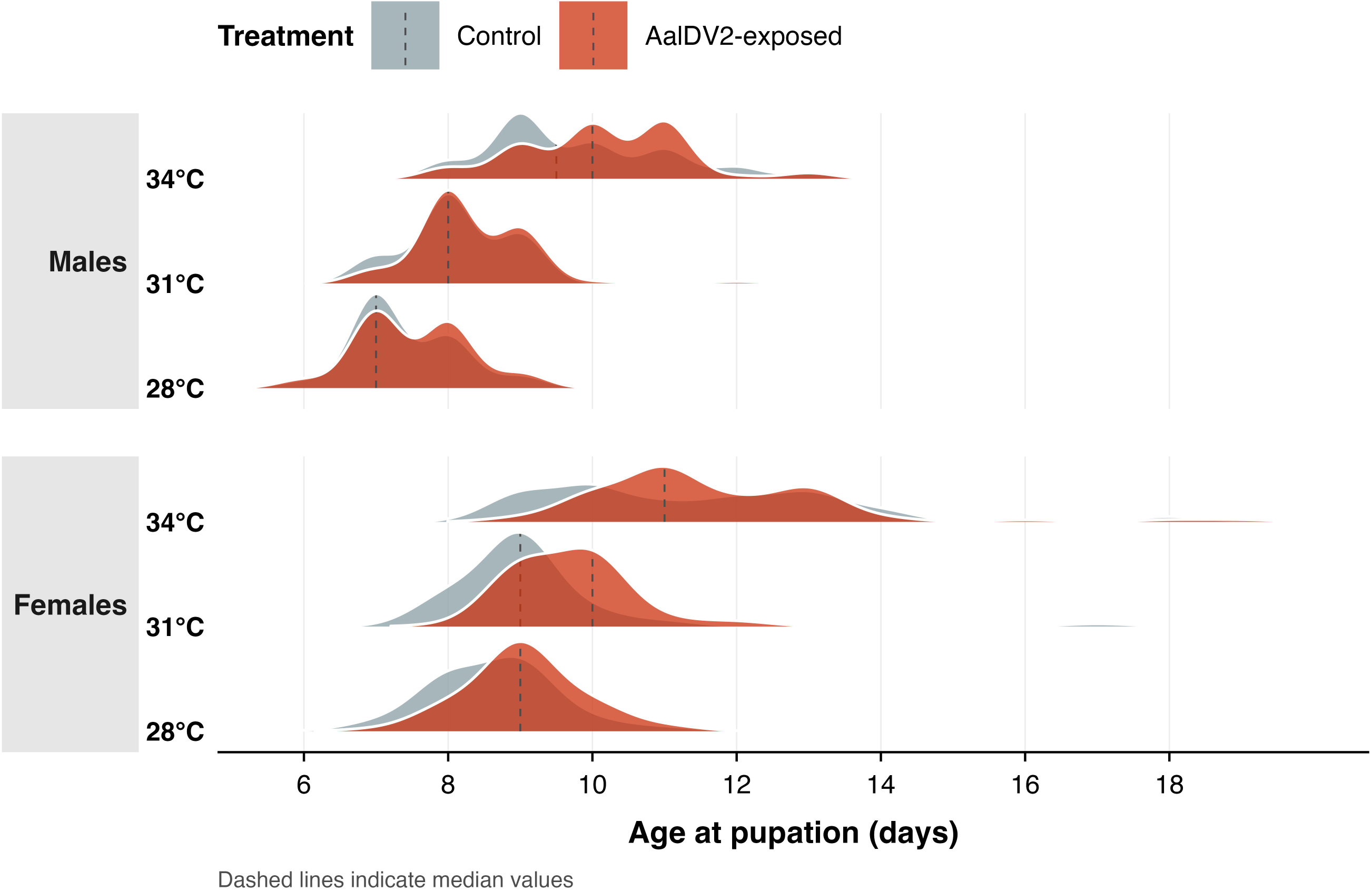
Distribution of the age at pupation (in days) for the mosquito *Aedes albopictus* exposed to different temperatures (28°C, 31°C, and 34°C) and to AalDV2. Data are stratified by sex and treatment (Control and AalDV2-exposed). Solid lines indicate the mean age at pupation, dashed lines indicate the median age at pupation. Colors represent different treatments: light blue for Control and dark red for AalDV2-exposed.

**Table 2.**
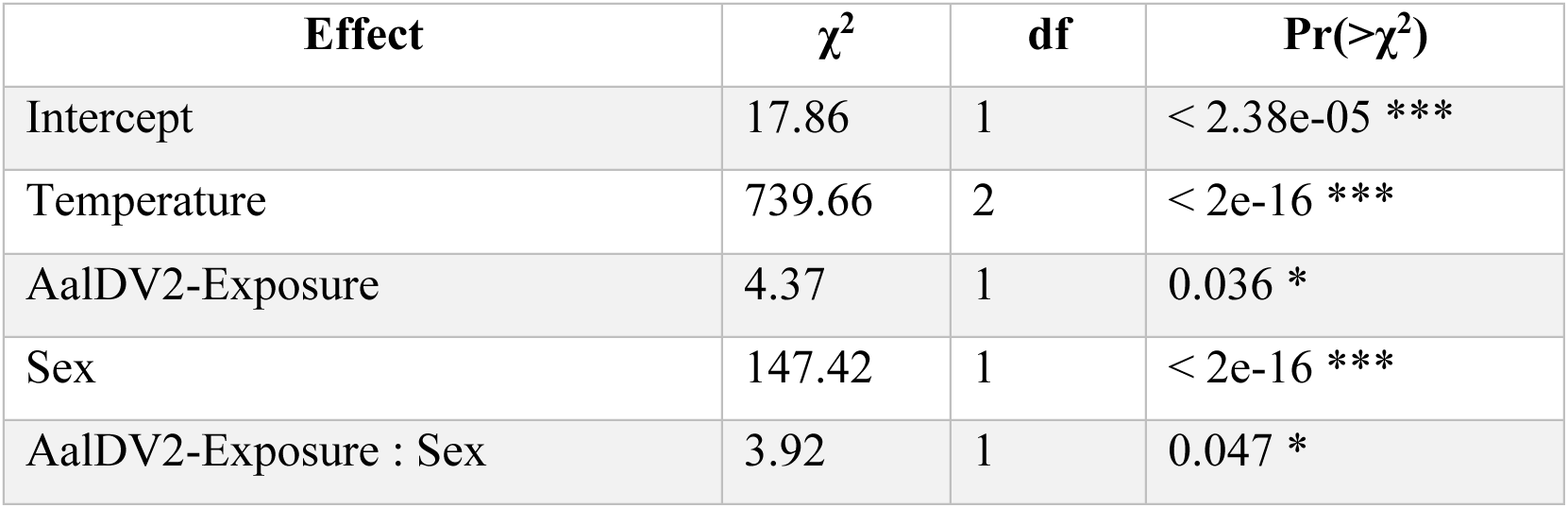
Analysis of the effect the temperature and the exposure to AalDV2 on the age at pupation. The response variable, age at pupation, was normalized using a double reversed Log_b (x + a) transformation to meet model assumptions. Fixed effects include Temperature, AalDV2-Exposure, Sex, and their interactions, with Bloc included as a random intercept. Chi-squared statistics (χ²), degrees of freedom (df), and p-values (Pr(>χ²)) are reported for each term. Significant effects are indicated as follows: *** (p < 0.001), ** (p < 0.01), * (p < 0.05).

| Effect | $\chi^2$ | df | Pr(>$\chi^2$) |
| --- | --- | --- | --- |
| Intercept | 17.86 | 1 | < 2.38e-05 *** |
| Temperature | 739.66 | 2 | < 2e-16 *** |
| AalDV2-Exposure | 4.37 | 1 | 0.036 * |
| Sex | 147.42 | 1 | < 2e-16 *** |
| AalDV2-Exposure : Sex | 3.92 | 1 | 0.047 * |

#### Impact on wing length

Adult wing length was used as a proxy for body size and was measured on 839 individuals for which we were able to measure the length of one or both wings. In the latter case, the arithmetic mean was taken as the wing size for a given mosquito. The analysis of wing size (Table 3) revealed a strong effect of sex (χ² = 1091.76, df = 1, p < 2.2e-16), with females having significantly larger wings than males across all temperatures, a difference that is more pronounced at lower temperature (Fig. 4). Temperature also had a significant effect (χ² = 23.14, df = 2, p = 9.44e-06), with individuals exposed to higher temperatures (31°C and 34°C) exhibiting reduced wing size as previously shown in *Anopheles gambiae s.s.* (Barreaux et *al.* 2018). Similarly, AalDV2-exposure had a significant negative effect (χ² = 9.71, df = 1, p = 0.0022), indicating that it reduces wing size. Interestingly, the effect of temperature on wing size was found to be sex-dependent and virus treatment-dependant, as indicated by the significant interaction between sex and temperature (χ² = 33.52, df = 2, p = 5.264e-08) and between temperature and treatment (χ² = 13.38, df = 2, p = 0.0012). Regarding temperature, it also appears that females exhibited a stronger decrease in wing size at higher temperatures compared to males.

**Fig. 4.**
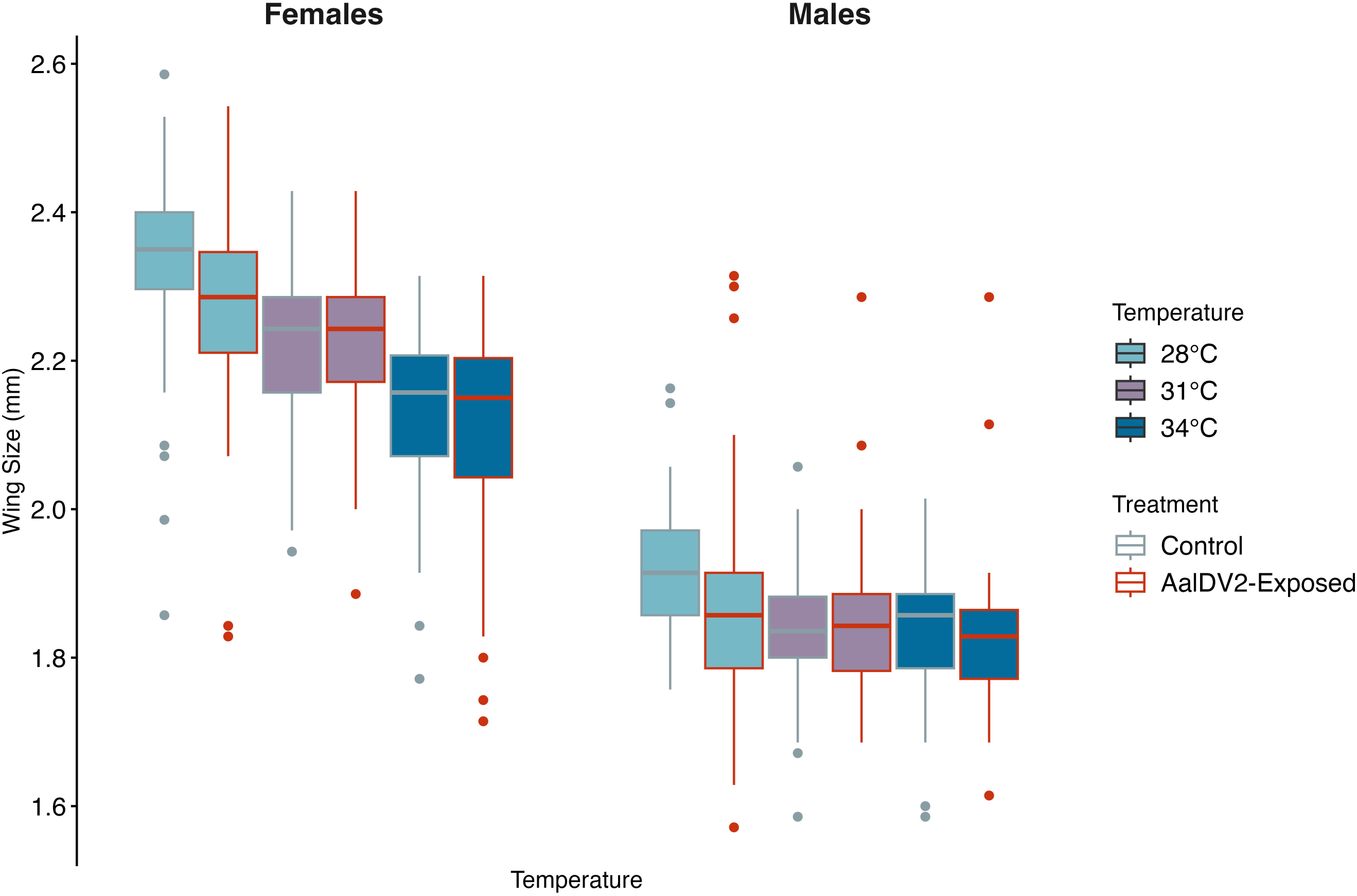
Distribution of wing size (in mm) in *Aedes albopictus* adults exposed to different temperatures (28°C, 31°C, and 34°C) and viral exposure (Control and AalDV2-exposed). Boxplots showing the Data are stratified by sex. Each boxplot represents the interquartile range (IQR) with the median indicated by the central line. Individual data points are overlaid to show the distribution of measurements. Colors represent different temperatures: light blue for 28°C, purple for 31°C, and dark blue for 34°C. Outlines distinguish treatments: grey for control and red for AalDV2-exposed.

**Table 3:** Analysis of the effect of temperature, sex and infection by the AalDV2 on the wing size of mosquitoes. The response variable, wing size, was transformed using the orderNorm transformation to ensure normality of residuals. Fixed effects in the model include Temperature, AalDV2-Exposure, sex, and their interactions, with Bloc included as a random intercept. Chi-squared statistics (χ²), degrees of freedom (df), and p-values (Pr(>χ²)) are reported for each term. Significant effects are denoted as follows: *** (p < 0.001), ** (p < 0.01), * (p < 0.05).

| Effect | $\chi^2$ | df | $\text{Pr}( > \chi^2 )$ |
| --- | --- | --- | --- |
| Intercept | 67.43 | 1 | <2.2 e-16 *** |
| AalDV2- Exposure | 9.71 | 1 | 0.0022 ** |
| Temperature | 23.14 | 2 | 9.44e-06*** |
| Sex | 1091.763 | 1 | <2.2 e-16 *** |
| Temperature : AalDV2- Exposure | 13.38 | 2 | 0.0012 *** |
| Sex : Temperature | 33.52 | 2 | 5.264e-08*** |

#### Impact on Fluctuating Asymmetry

The fluctuating asymmetry (FA) was analyzed on 478 individuals. It revealed a significant main effect of sex (χ² = 12.62, df = 1, p = 0.0004), with females exhibiting lower FA compared to males, indicating greater developmental stability (Fig. 5). No significant main effects of temperature (χ² = 3.40, df = 2, p = 0.18) or AalDV2 exposure (χ² = 0.04, df = 1, p = 0.85) were detected. However, a significant interaction between temperature and virus exposition treatment was observed (χ² = 6.08, df = 2, p = 0.048), suggesting that the effect of viral exposure on developmental stability depends on temperature.

**Fig. 5.**
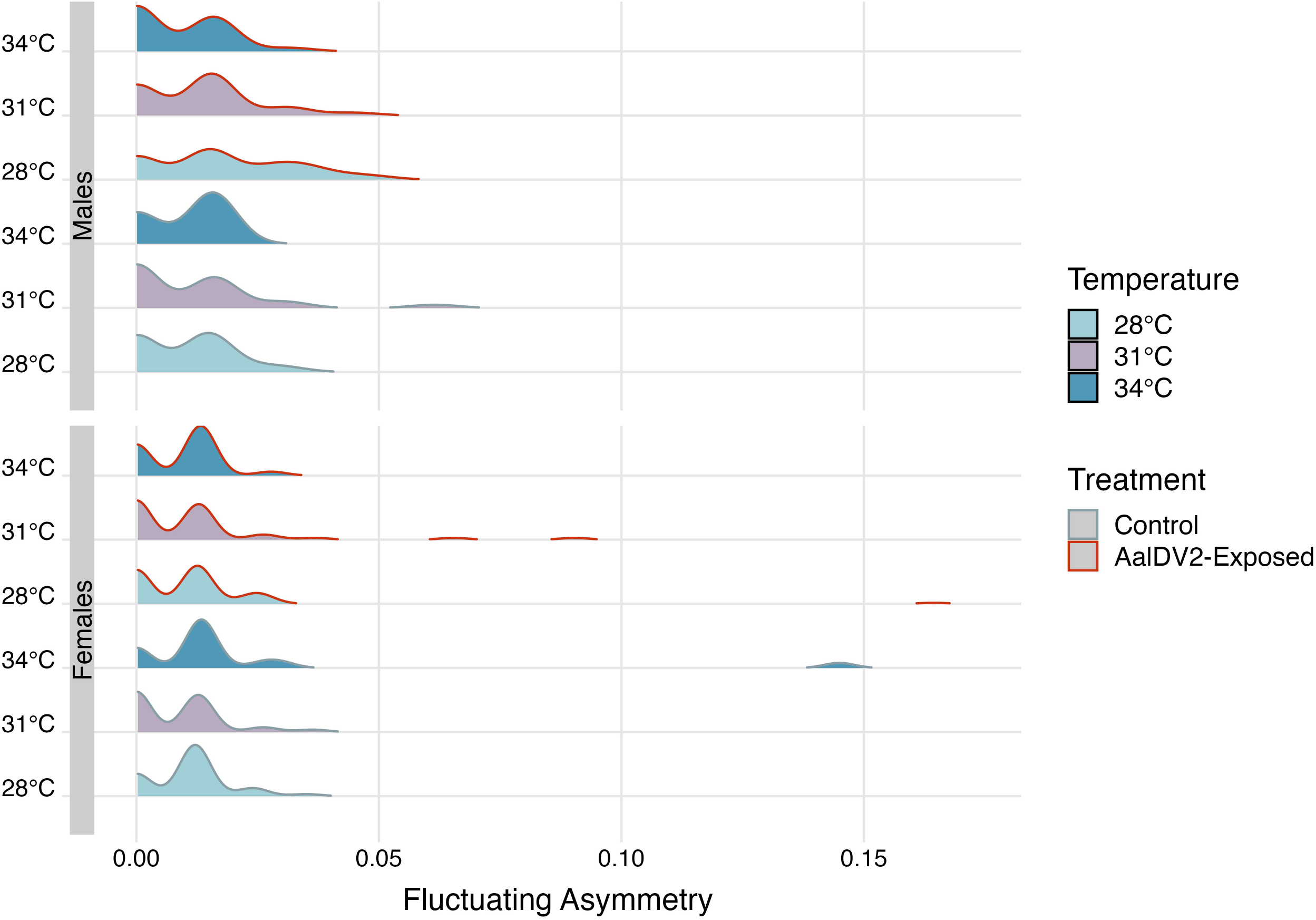
Distribution of the Fluctuating Asymmetry Index in *Aedes albopictus* exposed to different temperatures (28°C, 31°C, and 34°C) and viral exposure (Control and AalDV2-exposed). The FA Index is a size-scaled measure of asymmetry calculated as: FA Index = |Left Wing − Right Wing| / Mean (Left Wing, Right Wing). Data are stratified by sex. Colors represent different temperatures: light blue for 28°C, purple for 31°C, and dark blue for 34°C. Outlines distinguish treatments: gray for Control and red for AalDV2-exposed.

Post-hoc comparisons within each temperature showed that at 28°C, there was a trend toward higher FA in AalDV2-exposed individuals compared to controls (Estimate = -0.416, SE = 0.175, t = -2.373, p = 0.018). No significant differences were observed at 31°C (Estimate = 0.012, SE = 0.170, t = 0.069, p = 0.945). At 34°C, there was a non-significant trend toward lower FA in exposed individuals (Estimate = 0.260, SE = 0.145, t = 1.798, p = 0.073). The results suggest that the negative impact of viral exposure on developmental stability may be more pronounced under cooler conditions.

#### Effect of temperature on the infection prevalence and the viral load

Regarding the success of the infection, it has been split in 2 different parameters: the prevalence of the infection and the level of infection or viral load. It is noticeable that prevalence was particularly high, ranging from 96.27 to 100% and appears not to be temperature-dependent (Table S3). The analysis of the viral load revealed significant effects of temperature (F = 4.26, df = 2, p = 0.0148) and its interaction with sex (F = 5.39, df = 2, p = 0.0049) (Table 5; Fig. 6). This latter one revealed that temperature’s impact on viral load differed between males and females and post-hoc analyses confirmed that the effect of temperature was stronger in females (Estimate = 0.5310, p=0.0011).

**Fig. 6.**
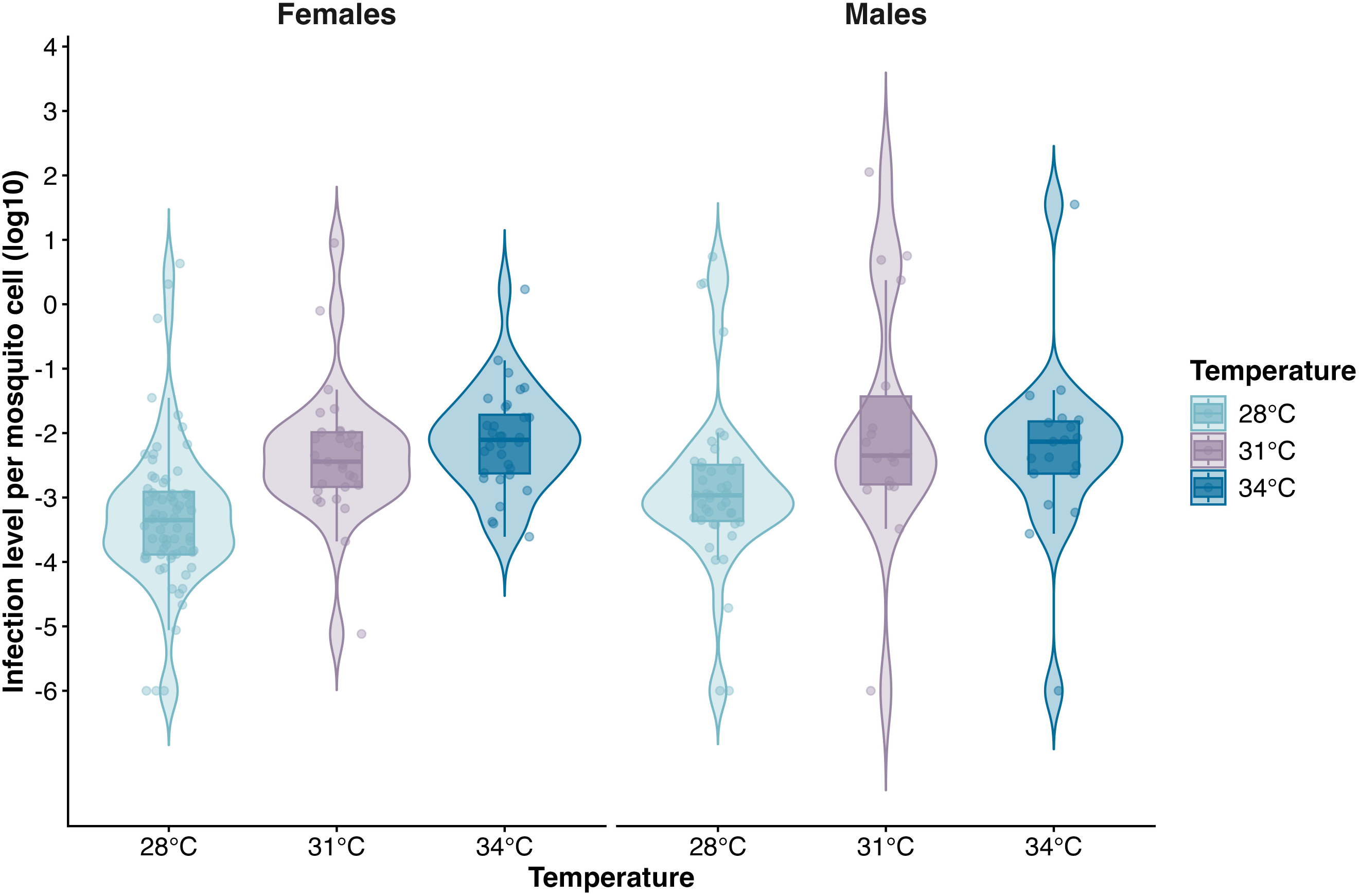
Distribution of AalDV2 infection levels (log₁₀-transformed viral RNA copies normalized to reference gene) in adult *Aedes albopictus* mosquitoes exposed to three temperature conditions (28°C, 31°C, and 34°C). Data are stratified by sex and each boxplot displays the median (center line), interquartile range (box), and 1.5× IQR range (whiskers). Individual data points are overlaid as jittered dots. Colors represent different temperatures: light blue for 28°C, purple for 31°C, and dark blue for 34°C.

**Table 4:** Analysis of the effect of temperature, sex and infection by the AalDV2 on the wing asymmetry of mosquitoes. The fluctuating asymmetry index was transformed using the orderNorm transformation to ensure normality of residuals. The fixed effects were Sex, Temperature, AalDV-exposure, and their interactions and Bloc was included as a random intercept. Chi-squared statistics (χ²), degrees of freedom (df), and p-values (Pr(>χ²)) are reported for each term. Significant effects are denoted as follows: *** (p < 0.001), ** (p < 0.01), * (p < 0.05), and (p < 0.1).

| Effect | $\chi^2$ | df | $\text{Pr}( > \chi^2 )$ |
| --- | --- | --- | --- |
| Intercept | 3.43 | 1 | 0.06 * |
| Sex | 12.62 | 1 | 0.000381 *** |
| Temperature | 3.4 | 2 | 0.18 |
| AalDV2-Exposure | 0.04 | 1 | 0.85 |
| AalDV2-Exposure : Temperature | 6.08 | 2 | 0.048 * |

**Table 5:**
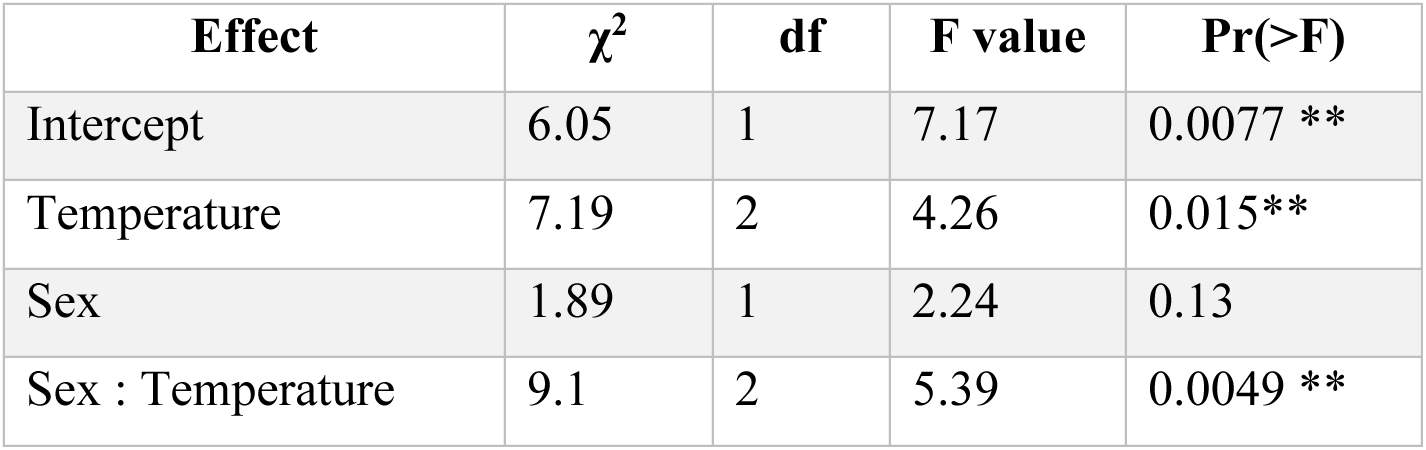
Analysis of the impact of the sex and temperature on the level of infection of *Aedes albopictus*. The table presents the chi-squared (χ²), degrees of freedom (df), F values, and p-values (Pr(>F)) for each effect. Significant effects are denoted as follows: *** (p < 0.001), ** (p < 0.01), * (p < 0.05), and (p < 0.1).

| Effect | $\chi^2$ | df | F value | Pr(>F) |
| --- | --- | --- | --- | --- |
| Intercept | 6.05 | 1 | 7.17 | 0.0077 ** |
| Temperature | 7.19 | 2 | 4.26 | 0.015** |
| Sex | 1.89 | 1 | 2.24 | 0.13 |
| Sex : Temperature | 9.1 | 2 | 5.39 | 0.0049 ** |

## Discussion

Our study highlights significant interactions between temperature and viral infection affecting a number of life-history traits in the tiger mosquito *Aedes albopictus*. This sheds light on how environmental factors may influence the effectiveness of densoviruses as a potential vector control tool. While our results complement previous studies highlighting negative impacts of densoviruses on mosquito life-history traits (Buchatsky 1989b, Carlson et *al.* 2006b, Li et *al*. 2019b, Perrin et *al.* 2020), they also reveal an unexpected protective beneficial effect of AalDV2 on mosquitoes at 34 °C. To our knowledge, this is the first evidence of such a positive, temperature-dependent impact of an entomopathogenic virus on mosquito survival. The heterogeneity observed between replicates at 34°C (∼10% difference) cannot be attributed to virus batch effects, as it occurred in both control and infected groups.

Regarding the effect of temperature on aquatic stages mortality, differences between 31 °C and 34 °C reveals a sharp nonlinear response, identifying a critical thermal threshold with potential implications. Beyond this threshold, virus-host interactions appear fundamentally altered, suggesting that biocontrol efficacy may be highly context-dependent and hardy predictable in warming climates. Linear extrapolations are therefore insufficient for modeling mosquito population responses to rising temperatures (Ewing et *al*. 2016) as well as when evaluating potential biological control agents. This protective effect suggests viral modulation of physiological stress responses, indicating that host-virus-environment interactions are more complex than simple additive effects, as previously shown in honey bees (Hsieh and Dolezal 2024). Even if they appear counterintuitive, similar phenomena have already been documented in other systems. Viruses can indeed confer thermal tolerance to their hosts, as seen with fungal endophyte-virus-grass interactions (Marquéz et *al*. 2007), with the Tomato yellow leaf curl virus (TYLCV) improving plant survival under heat and drought (Gorovits et *al*. 2022), and RNA viruses improving plant tolerance to freezing (Xu et *al*. 2008, Westwood et *al*. 2013). In aquatic mollusks, high trematode infection intensities have nullified temperature impacts on blue mussels (Selbach et *al*. 2020). In insects, thermal stress acting as immune priming has increased survival of infected *Galleria mellonella* larvae (Browne et *al*. 2014) and thermal challenge duration affects the outcomes of the infection by the cell-fusing agent virus (CFAV) in *A. albopictus* (Perdomo et *al*. 2025). A speculative mechanism could involve heat shock protein (HSP) upregulation induced by multiple stressors including infections and heat (Feder and Hofmann 1999).

Regarding the development, exposure to AalDV2 lead to a sex-dependent delay, with females experiencing greater costs than males. This could have population-level implications with extended aquatic periods in infected females increasing vulnerability to environmental hazards, predation, and resource competition. While potentially affecting generation time and slowing population growth, making this effect favorable for vector control, this could also drive sex-specific evolutionary responses to infection (Restif and Amos 2010). Critically, whether these developmental delays affect adult longevity, fecundity, and vector competence remains unknown and is essential for predicting biocontrol efficacy.

Regarding wing size, the dual impact of temperature and viral infection, with females showing greater temperature sensitivity, confirms that both genetic and environmental factors shape this fitness-related trait (Christiansen-Jucht et *al*. 2015) that is correlated with fecundity (Armbruster and Hutchinson 2002). Concerning the fluctuating asymmetry of wings, the marginally significant interaction between temperature and AalDV2 exposure suggests potentially opposing effects of viral infection on developmental stability depending on thermal conditions. The trend toward destabilization at 28 °C but stabilization at 34 °C parallels the survival patterns, hinting at a shift in virus-host interactions across temperatures. However, these trends require validation to determine functional consequences for mosquito fitness and vector competence. One plausible explanation for the observed pattern at 34 °C could be a survival bias. Higher temperatures may impose stronger selective pressures, leading to the survival of only those individuals with lower FA. If individuals with higher FA did not survive the aquatic stage at 34 °C, the remaining population would predominantly consist of those with lower FA, potentially masking the true effect of AalDV2 exposure on developmental stability at this temperature. Future studies could investigate survival rates in relation to FA at different temperatures to test this hypothesis directly.

When considering the AalDV2-infection itself, the consistently high prevalence (>96%) across all temperatures represents a critical biocontrol characteristic, ensuring reliable performance under variable field conditions. The contrast between the temperature-independent prevalence and the temperature-dependent viral load reveals that the initial infection is qualitatively robust, but replication dynamics are thermally sensitive leading to quantitative differences. The sex-dependent increase in viral load with temperature warrants careful consideration. Several mechanisms may explain this pattern such as a longer female developmental time, especially at high temperatures, providing extended replication windows as well as differences in immune responses between sexes. For vertical transmission, a potential transmission route for densoviruses (Altinli et *al.* 2018), higher female viral loads could enhance transovarial transmission efficiency, a critical point for densovirus persistence. Understanding these sex-specific dynamics appears essential for predicting ecological outcomes and evolutionary responses in vector control programs.

From a transmission ecology perspective with vector control in mind, high infection rates facilitate both horizontal transmission through contaminated breeding sites and vertical transmission across generations. However, this is double-edged: while favorable for establishing viral presence, the protective effect at extreme temperatures could allow infected mosquitoes to persist under otherwise lethal conditions, potentially enhancing mosquito resilience in warming climates.

These results underscore the necessity of considering temperature-specific effects when evaluating biological control agents and have critical implications for vector control in a warming world. The complexity of the interactions between an environmental stressor, a viral infection and host physiology can produce unexpected outcomes potentially difficult to predict under field conditions. This is especially important given observations of densovirus symbiosis in other systems (Xu et *al.* 2014) and highlights the interest to consider mosquito viruses as endosymbionts (Altinli et *al.* 2020). Whether virus-mediated thermal tolerance persists in adults is critical, as this could affect not only population dynamics but also arbovirus transmission capacity. Current studies on densovirus impacts on arbovirus transmission remain limited, use artificial systems, and show conflicting results (Berger et *al.* 2025). While AalDV2’s high infection success and negative fitness impacts suggest biocontrol potential, our findings raise critical questions about ecological and evolutionary implications. If AalDV2 enhances larval survival at extreme temperatures, deployment could inadvertently strengthen mosquito resilience in warming climates, potentially undermining control efforts in regions most affected by neglected tropical diseases.

Overall, our study reveals a paradoxical outcome in the AalDV2-*Aedes albopictus* interaction: while the virus imposes fitness costs through delayed development and reduced body size, it confers an unexpected survival advantage at extreme temperatures. This represents the first documentation of a protective effect by an entomopathogenic virus on mosquito survival under thermal stress, challenging conventional assumptions about pathogen impacts and highlighting the context-dependency of host-parasite interactions.

Moving forward, research should prioritize several key areas such as the elucidation of the physiological mechanisms underlying virus-mediated thermal tolerance and the evaluation of any protective effects persisting through the adult stage and potentially influencing the vector competence. A step further could then permit to evaluating the long-term population and epidemiological consequences under realistic field conditions with natural temperature fluctuations. Only through such comprehensive investigation can we determine whether densoviruses represent a viable tool for *Aedes albopictus* control in the context of climate change. These findings underscore a fundamental principle: effective biocontrol strategies must account for the full complexity of host-parasite-environment interactions rather than relying on simplified predictions of pathogen effects.

## Conclusion

The changes of temperatures due to climate changes remains uncertain and its impact on mosquito populations dynamics is difficult to predict. Understanding how such changes will affect the life-history traits of mosquitoes and especially invasive species such as *Aedes albopictus* is a major concern to determine their influence on the vector populations dynamics and to forecast their potential geographic spread and the associated risk of disease transmission. The development of novel tools for vector control should also take into account the impact of temperature on their efficacy in a variety of contexts. The effects of temperature on *Aedes albopictus* life-history traits, particularly in the context of AalDV2 infection, highlight the complexity of host-parasite-environment interactions. While AalDV2 infection appears to provide a survival advantage at high temperatures, this effect raises important questions about the potential ecological and evolutionary consequences of using this virus in vector control. The strong temperature-dependent effects on survival, sex-ratio, developmental timing, and infection levels underscore the need to consider environmental variables when developing novel vector control strategies. As global temperatures continue to rise, understanding how these interactions will evolve is crucial for forecasting the spread of *Aedes albopictus* and the associated risk of arbovirus transmission. These findings also emphasize the importance of considering both biological and environmental factors in designing and implementing effective biocontrol interventions, particularly in the face of climate change.

## Acknowledgements

We are grateful to Doriane Mutuel for the maintenance of the cell cultures, to Hélène Sobry for the production of the AalDV2 virus, to Philippe Clair from the qPCR platform at Montpellier University, to Aurélie Perrin for discussions and to Hélène Sobry and Pierre-Alain Girard for technical assistance. This work was funded by the French ANSES Environment-Santé-Travail research programme (PNR EST) (project 2018/1/183, “DENSOTOOL”, 2019–2023).

