## Supplementary Information for "Stranger Swings: Temperature-Dependent Upsides and Downsides of a Densovirus in *Aedes albopictus*"

| Bloc | Temperature | Treatment | Total larvae | Dead Larvae | Dead pupae | Dead adult at emergence | Total Death | Surviving after emergence | Percentage of surviving mosquitoes | IntConf_low of the Percentage of surviving mosquitoes | IntConf_sup of the Percentage of surviving mosquitoes |
|---|---|---|---|---|---|---|---|---|---|---|---|
| 1 | 28 | C | 47 | 1 | 4 | 0 | 5 | 42 | 89.36 | 77.40 | 95.37 |
| 1 | 28 | If | 95 | 2 | 2 | 0 | 4 | 91 | 95.79 | 89.67 | 98.35 |
| 1 | 31 | C | 35 | 2 | 1 | 0 | 3 | 32 | 91.43 | 77,62 | 97.04 |
| 1 | 31 | If | 92 | 5 | 1 | 2 | 8 | 84 | 91.30 | 84.77 | 95.53 |
| 2 | 28 | C | 95 | 0 | 2 | 0 | 2 | 93 | 97.89 | 92,65 | 99.42 |
| 2 | 28 | If | 186 | 5 | 4 | 0 | 9 | 177 | 95.16 | 91.06 | 97.43 |
| 2 | 34 | C | 88 | 24 | 25 | 0 | 49 | 39 | 44.32 | 34.39 | 54.72 |
| 2 | 34 | If | 181 | 55 | 30 | 0 | 85 | 96 | 53.04 | 45.78 | 60.17 |
| 3 | 31 | C | 96 | 5 | 4 | 0 | 9 | 87 | 90.62 | 90.62 | 83.13 |
| 3 | 31 | If | 95 | 8 | 3 | 0 | 11 | 84 | 88.42 | 88.42 | 80.45 |
| 3 | 34 | C | 158 | 99 | 44 | 0 | 143 | 15 | 9.49 | 5.84 | 15.07 |
| 3 | 34 | If | 155 | 83 | 37 | 3 | 123 | 32 | 20.65 | 15.02 | 27.69 |

**SI Table S1**: Survival of the mosquitoes up to emergence under different conditions. The table presents the survival data of mosquitoes from larval to adult stages under varying temperatures and exposure to the AalDV2 virus across different blocks. The percentage of surviving mosquitoes and its 95% confidence intervals (CI) were calculated with the *binconf* function from the *Hmisc* package in R.

| Effect | $\chi^2$ | df | Pr(>$\chi^2$) |
|---|---|---|---|
| Intercept | 0.50 | 1 | 0.48 |
| AalDV2 - Exposure | 0.87 | 1 | 0.35 |
| Temperature | 1.12 | 2 | 0.57 |

**SI Table S2**: Statistical analysis of the effect the temperature and the exposure to AalDV2 on the sex-ratio at emergence. The response variable is the proportion of females at emergence, modeled using a binomial generalized linear mixed model (GLMM) with a logit link function. Degrees of freedom (df), chi-squared statistics ($\chi^2$), and p-values (Pr(>$\chi^2$)) are reported for each fixed effect. None of the tested effects (AalDV2 treatment or temperature) significantly influenced the sex ratio at emergence ($p > 0.05$).

| Sex | Temperature | Number of mosquitoes surviving emergence after exposure to AalDV2 | Number of non-infected at adulthood | Percentage infected | IntConf_low | IntConf_sup |
|---|---|---|---|---|---|---|
| Male | 28 | 134 | 3 | 97.76 | 97.76 | 93.62 |
| Male | 31 | 84 | 1 | 98.81 | 98.81 | 93.56 |
| Male | 34 | 58 | 1 | 98.28 | 98.28 | 90.86 |
| Female | 28 | 134 | 5 | 96.27 | 96.27 | 91.56 |
| Female | 31 | 84 | 1 | 98.81 | 98.81 | 93.56 |
| Female | 34 | 70 | 0 | 100 | 100 | 94.8 |

**SI Table S3**: **Percentage of AalDV2 infection in adult mosquitoes by sex and temperature.** This table presents the infection status of mosquitoes that survived to adulthood after exposure to the AalDV2 virus, categorized by sex and temperature. The percentage of infected mosquitoes and its 95% confidence intervals (CI) were calculated with the binconf function from the Hmisc package in R.
